# Lysosome Exocytosis Inhibition Impairs Triple-Negative Breast Cancer Cell Invasion

**DOI:** 10.64898/2026.11.20.765315

**Authors:** Ana R. Rodrigues, Cristina Escrevente, Teresa Barona, Inês Fonseca, Mariana Feliciano, Matilde G. Oliveira, José S. Ramalho, Duarte C. Barral

## Abstract

Breast cancer (BC) is the most frequently diagnosed cancer in women, with metastasis remaining the leading cause of mortality. Among BC molecular subtypes, triple-negative BC (TNBC) is the most aggressive, characterized by high metastatic potential and limited treatment options, underscoring the need for developing new therapeutic strategies. Cancer progression relies on the subversion of multiple cellular processes, among which lysosome exocytosis is described as a major driver of cell invasion. Indeed, lysosome exocytosis has been shown to play a role in extracellular matrix degradation and remodeling, which is particularly relevant to sustain cancer cell invasive capacity. In this study, we show that highly-invasive BC cell lines present increased levels of lysosome exocytosis when compared to non- or low-invasive ones. To investigate the functional effect of inhibiting this pathway on TNBC cell invasion, we used a genetic approach, by silencing RAB10 or a pharmacological strategy, using rosmarinic acid, and assessed invasive capacity. Both approaches significantly decreased lysosome exocytosis, measured as the release of the lysosome enzyme β-hexosaminidase, and strongly impaired TNBC cell invasive capacity. Together, these findings highlight the potential of lysosome exocytosis as a therapeutic target to impair TNBC cell invasion, hindering BC progression, and improving patient prognosis.

## Introduction

Breast cancer (BC) is one of the most prevalent malignancies worldwide, with an estimated 2.3 million new cases diagnosed in 2022, according to the World Health Organization ^1^. Despite advances in early detection and therapeutic strategies, BC remains a leading cause of cancer-related mortality in women, being responsible for approximately 670,000 deaths in 2022 ^1–3^. Among its different molecular subtypes, triple-negative BC (TNBC), characterized by lack of estrogen receptor (ER), progesterone receptor (PR), and human epidermal growth factor 2 (HER2) expression, is known by its invasive capacity, lack of efficient targeted therapies, high metastatic potential, and poor clinical prognosis ^2,4^. Since metastasis is the major cause of cancer-related deaths, there is an urgent need to develop new strategies that target this process to improve patient outcomes.

Metastatic dissemination requires that cells acquire migratory and invasive capacities, allowing them to escape from the primary tumor site and enabling colonization of distant organs ^5^. For this, cancer cells modulate multiple intracellular mechanisms, among which lysosome exocytosis has emerged as a key contributor ^6,7^. Through this process, lysosomes fuse with the plasma membrane (PM) in a Ca^2+^-dependent manner, and release hydrolytic enzymes such as cathepsins to the extracellular space ^8^. This promotes extracellular matrix (ECM) degradation and local acidification of the tumor microenvironment (TME), providing an optimal environment for cancer cell migration and invasion ^6,7,9^. Moreover, lysosome exocytosis supports tumor progression through multiple pro-tumorigenic processes. Specifically, it promotes chemoresistance by facilitating the efflux of lysosomotropic chemotherapeutic agents, promotes the activation of epithelial-to-mesenchymal transition, supports angiogenic signaling, and contributes to immune evasion ^9–11^. In addition, lysosome exocytosis enables PM repair and drives ECM remodeling and degradation^6,7^. Moreover, lysosome exocytosis was shown to be upregulated and to correlate with increased cell invasive capacity in different cancer types, namely, sarcoma, melanoma, and glioma ^11–14^. Furthermore, lysosome exocytosis was identified as a driver of tumor progression, including in BC radiation-surviving cancer cells ^15^. In this context, targeting lysosome exocytosis could be a promising strategy to impair TNBC cell migration and invasion with a reduced risk of secondary effects resulting from general lysosomal inhibition.

Lysosome exocytosis is tightly regulated by a coordinated molecular machinery. Lysosome fusion with the PM is mediated by the interaction of the Ca^2+^ sensor Synaptotagmin VII ^16^ with the SNARE complex composed by VAMP7, Syntaxin-4 and SNAP-23 ^17^. Moreover, our group showed that the Rab small GTPases RAB11A and RAB11B transiently interact with lysosomes at the cell periphery ^18^ and that RAB3A is essential for lysosome positioning before fusion ^19^. Additionally, the silencing of RAB10, a small GTPase implicated in multiple cellular functions including polarized exocytosis ^20^ and early-to-recycling endosome maturation ^21^, was shown to impair lysosome exocytosis and PM repair in THP-1 and HeLa cell lines ^19,22^. Therefore, targeting these molecular players is an attractive approach to modulate lysosome exocytosis and evaluate the relevance of lysosome exocytosis in tumor progression. Besides genetic approaches, the use of small-molecule modulators offers a pharmacologically relevant strategy to target lysosome exocytosis. Indeed, a few compounds have been reported to interfere with this pathway, including verapamil, a Ca^2+^ channel blocker, which was reported to impair lysosome exocytosis and drug efflux in sarcoma cells ^6^; and vacuolin-1, a potent inhibitor of PIKfyve – required for lysosome maturation, shown to inhibit lysosome exocytosis in HeLa cells ^23,24^.

Although lysosome exocytosis has been implicated in cancer cell invasion, it remains to be shown whether its targeting is sufficient to impair cancer cell invasive capacity. Here, we report that highly invasive BC cell lines present increased levels of lysosome exocytosis when compared with non-invasive or low-invasive BC cell lines. Considering this observation, we investigated whether inhibition of lysosome exocytosis impairs TNBC cell invasive capacity, using both genetic and pharmacological approaches. Our results demonstrate that the inhibition of lysosome exocytosis effectively impairs TNBC cell invasion. These findings provide evidence that targeting lysosome exocytosis represents a promising therapeutic strategy to be used alone or in combination with current therapies to delay TNBC progression and increase patients’ survival rate.

## Results

### Lysosome exocytosis is enhanced in triple-negative breast cancer cells

To determine whether lysosome exocytosis correlates with BC cell invasion, we compared lysosome exocytosis levels across BC cell lines with different invasive capacities. For this, we used MCF-7 luminal A cell line, which are low-invasive; MDA-MB-231 and HCC1806 TNBC cell lines, described to be highly invasive; and MCF10A as a non-tumorigenic, non-invasive cell line. We analyzed lysosome exocytosis by measuring the levels of the late endosome/lysosome marker LAMP1 at the cell surface by flow cytometry, and excluding dead cells by propidium iodide (PI) staining. Under constitutive conditions, highly invasive MDA-MB-231 cells were found to display significantly higher LAMP1 surface levels compared with both the low-invasive MCF-7 and non-invasive MCF10A cells (Fig. 1a). Similarly, highly invasive HCC1806 cells were also found to display significantly increased LAMP1 surface levels compared to the low-invasive MCF-7 cells (Fig. 1a). Given that lysosome exocytosis occurs constitutively at low levels, but can be enhanced by increased Ca^2+^ levels, we used the Ca^2+^ ionophore ionomycin to stimulate lysosome exocytosis prior to LAMP1 surface level quantification. Under these conditions, the results revealed an even more striking increase in lysosome exocytosis levels in the highly invasive cell lines, with both MDA-MB-231 and HCC1806 cells exhibiting significantly higher LAMP1 surface levels, compared with MCF10A and MCF-7 cells (Fig. 1b). To further confirm these results and considering that LAMP1 is not only present in lysosomes but also in late endosomes, we measured the ionomycin-stimulated release of the lysosomal hydrolytic enzyme β-hexosaminidase to the culture medium, which is a more specific readout of lysosome exocytosis. Consistent with the LAMP1 results, we observed that highly invasive cell lines present increased β-hexosaminidase secretion compared with the low- and non-invasive cell lines (Fig. 1c), further supporting that lysosome exocytosis is increased in TNBC cells.

**Figure 1.**
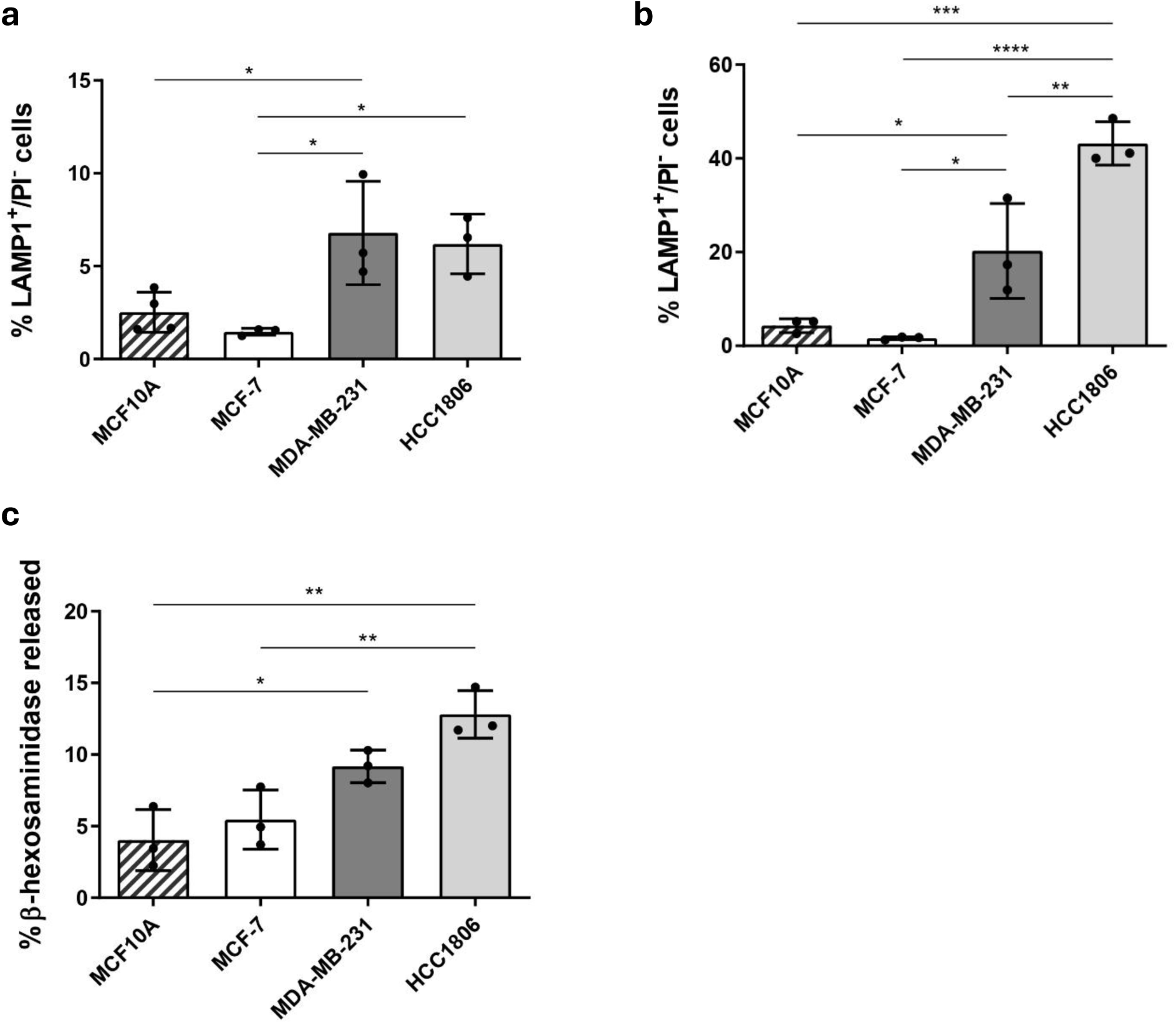
Lysosome exocytosis levels are enhanced in highly invasive triple-negative breast cancer cells. MCF10A, MCF-7, MDA-MB-231 and HCC1086 cells were **(a)** stained with Alexa Fluor 488 anti-LAMP1 antibody and analyzed by flow cytometry to assess cell surface LAMP1 levels under constitutive conditions or **(b)** stimulated with 1 μM ionomycin and 4 mM CaCl_2_ for 5 min. at 37 °C to induce Ca^2+^-dependent lysosome exocytosis, and stained with Alexa Fluor 488 anti-LAMP1 antibody, and analyzed in the same manner. The percentage of LAMP1-positive (LAMP1+) cells and propidium iodide-negative (PI−) cells was quantified. **(c)** β-hexosaminidase release was measured in cell extracts and supernatants and quantified as described in the Materials and Methods. Results represent the mean ± s.d. of three independent experiments. Statistical significance was determined using one-way ANOVA followed by Dunnett’s multiple comparisons test (****P<0.0001, ***P<0.001, ** for P<0.01, *P<0.05).

### Rab10 silencing impairs lysosome exocytosis and triple-negative breast cancer cell invasion

To ascertain that the modulation of lysosome exocytosis affects TNBC cell invasion, we started by investigating the effect of silencing Rab10 in lysosome exocytosis and TNBC cell invasion. RAB10 was selected as a candidate regulator because it has been previously implicated in lysosome exocytosis in THP-1 and HeLa cells ^19,22^; however, its role in regulating lysosome exocytosis has not been demonstrated in TNBC cells. For this, we used shRNA-expressing lentiviruses to silence RAB10 (shRAB10 F6 and F9) in MDA-MB-231 cells. Cells transduced with non-targeting shRNAs (Mission) were used as a control. Silencing efficiency was confirmed by RT-PCR (Supplementary Fig. S1). Lysosome exocytosis was assessed upon stimulation with ionomycin, by measuring β-hexosaminidase secretion. We observed that RAB10 silencing leads to a significant decrease in lysosome exocytosis for both shRNA hairpins tested (Fig. 2a). Next, to assess the effect of lysosome exocytosis inhibition, triggered by RAB10 silencing in TNBC cell invasion, we performed Matrigel-coated transwell invasion assays. We found that RAB10-silenced cells display a significant decrease in the invasion capacity, when compared to the control (Fig. 2b, c). To further investigate the effects of RAB10 silencing on TNBC cell invasion, a gelatin degradation assay was performed, as a complementary approach to the Matrigel-coated transwell invasion assay. For this, the gelatin degradation area was quantified as the ratio of the image area with gelatin degradation (black spots), over the area of the entire image. The results showed a reduction in the degradative capacity of RAB10-silenced cells when compared with control cells, although not statistically significant (Fig. 2d, e). The percentage of cells with active invadopodia, *i.e.* cells presenting colocalization between cortactin staining and gelatin degradation spots, was also determined. However, no significant difference between RAB10-silenced cells and the vehicle control (PBS) was observed (Fig. 2f). Overall, these observations confirm that RAB10 is a positive regulator of lysosome exocytosis in TNBC cells and show that RAB10 silencing impairs the invasive potential of MDA-MB-231 TNBC cells, possibly by decreasing ECM degradation.

**Figure 2.**
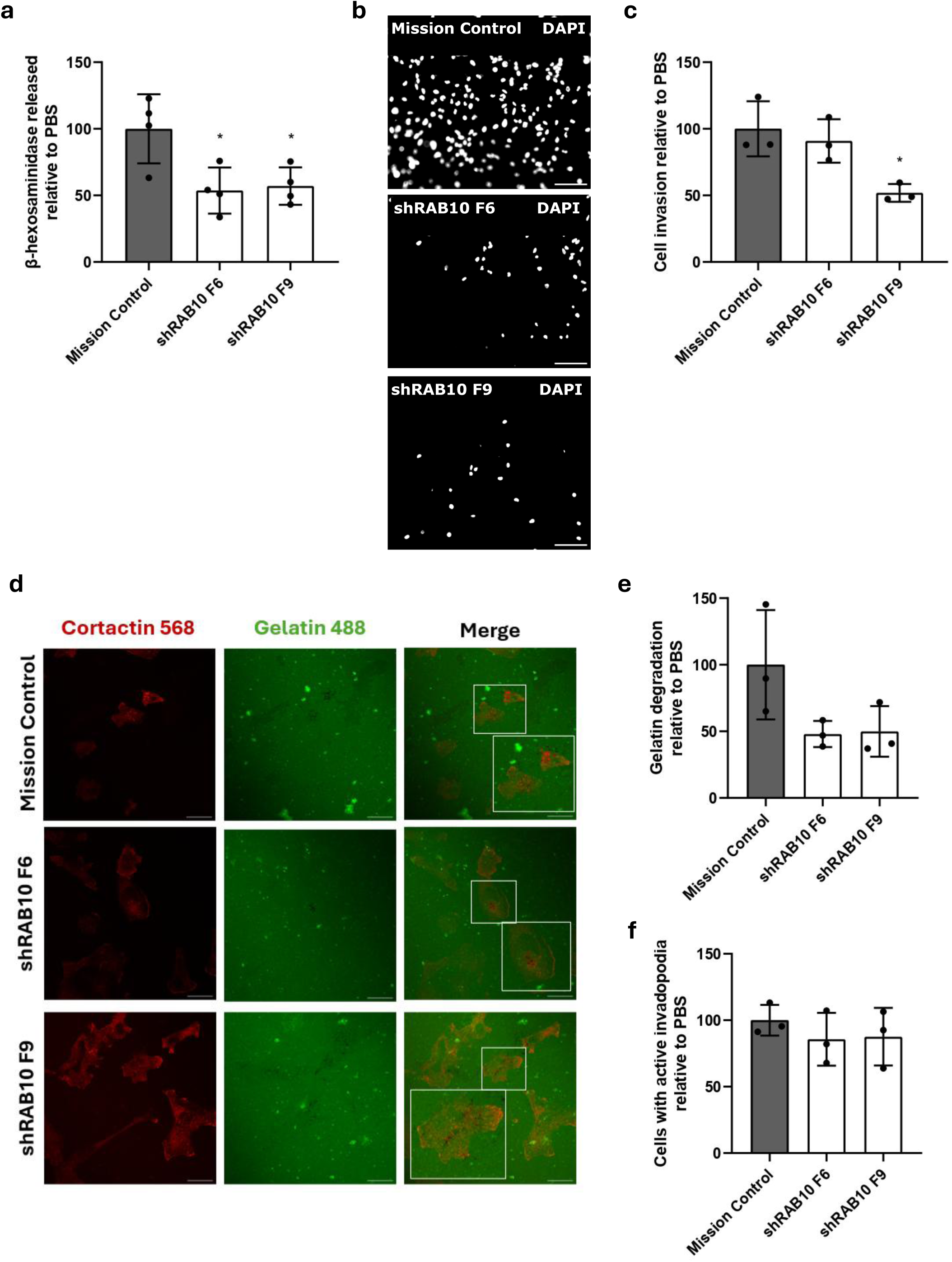
RAB10 silencing decreases lysosome exocytosis and impairs invasion of triple-negative breast cancer cells. MDA-MB-231 cells were transduced with lentiviruses encoding shRNA hairpins targeting RAB10 (shRAB10 F6 and F9), as well as a non-targeting shRNA (Mission), as a control. **(a)** Cells were stimulated with 1 μM ionomycin and 4 mM CaCl_2_ for 5 min. at 37 °C to trigger Ca^2+^-dependent lysosome exocytosis and cell extracts and supernatants were collected. β-hexosaminidase release was quantified as described in the Materials and Methods. **(b)** To measure invasion, cells were seeded into the upper chamber of Matrigel-coated transwells in serum-free media. After 24 h, invading cells were fixed and stained with DAPI. Scale bar = 100 µm. **(c)** Quantification was performed by counting DAPI-stained cells in 10 randomly-selected fields per insert. Values were normalized to the control, set at 100%. **(d)** Representative confocal images of cells seeded on coverslips coated with Oregon Green 488 gelatin (green) and stained with mouse anti-cortactin (red). Scale bar = 20 μm. **(e)** Percentage of cells with active invadopodia. **(f)** Percentage of gelatin degradation area. Results are represented as mean ± s.d. of at least three independent experiments. Statistical significance was determined using one-way ANOVA with Dunnett’s multiple comparison test (*P<0.05).

### Inhibition of lysosome exocytosis using rosmarinic acid impairs lysosome exocytosis and triple-negative breast cancer cell invasion

To evaluate whether pharmacological inhibition of lysosome exocytosis affects TNBC cell invasion, we first explored known inhibitors, namely, verapamil and vacuolin-1 ^6,23,24^. However, we were unable to observe any significant changes in lysosome exocytosis upon incubation of MDA-MB-231 cells with either one of these compounds (Supplementary Fig. S2a and S2b), suggesting their effect is likely cell-specific and highlighting the need to identify drugs that effectively impair lysosome exocytosis in TNBC cells. Phenolic compounds are promising anti-cancer agents ^25^. In particular, rosmarinic acid (RA), a phenolic acid derived from rosemary, was proposed as a negative modulator of purinergic receptors, including P2X7, a Ca^2+^-permeable ion channel ^26^. Therefore, we hypothesized that RA-mediated inhibition of P2X7 receptor suppresses the levels of intracellular Ca^2+^, which is required for lysosome-PM fusion, consequently impairing lysosome exocytosis ^26,27^. To test this, MDA-MB-231 cells were incubated with RA at different concentrations (0.07, 0.14, 0.27, 0.56, 0.83 and 1.11 mM), for 24 h. As a control, we used the vehicle (PBS). After 24 h, lysosome exocytosis was quantified by measuring the release of β-hexosaminidase to the culture medium. The results revealed that RA effectively decreases lysosome exocytosis in a concentration-dependent manner, consistent with our hypothesis (Fig. 3a). A trypan blue exclusion assay confirmed that the RA concentrations used did not affect MDA-MB-231 cell proliferation, with exception of 1.11 mM that showed a significant reduction in cell number 24 h after RA incubation (Fig. S3). Our results were further confirmed using a resazurin viability assay. Indeed, incubation of MDA-MB-231 cells with RA concentrations bellow 1.11 mM for 24 h did not affect cell viability (Fig. 3b). Therefore, the concentration of 0.83 mM was selected for subsequent experiments. We next assessed whether RA-mediated inhibition of lysosome exocytosis translates into impaired cell invasive capacity. For this, MDA-MB-231 cells were pre-incubated with 0.83 mM of RA for 2 h, seeded onto Matrigel-coated transwells, and allowed to invade for 24 h in the presence of RA. We observed that MDA-MB-231 cell invasion decreases significantly in RA-incubated cells, when compared to the control (Fig. 3 c, d). Together, these results demonstrate that pharmacological inhibition of lysosome exocytosis is sufficient to impair TNBC cell invasive behavior.

**Figure 3.**
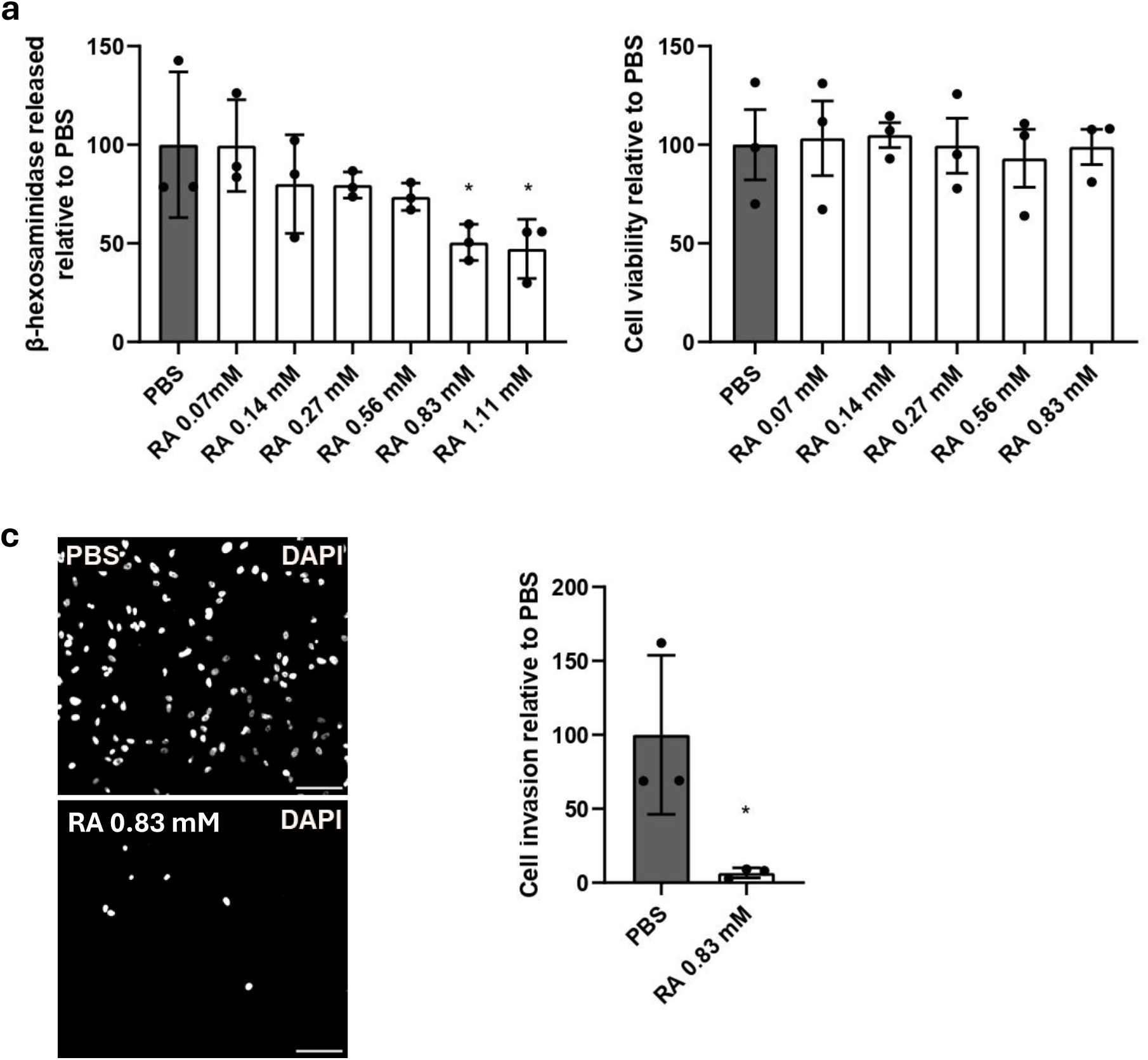
Rosmarinic acid inhibits lysosome exocytosis and impairs triple-negative breast cancer cell invasion. MDA-MB-231 cells were incubated with RA at different concentrations (0.07, 0.14, 0.27, 0.56, 0.83 and 1.11 mM) for 24 h, with PBS as the vehicle control. **(a)** β-hexosaminidase release was measured in cell supernatants and in extracts and quantified as described in the Materials and Methods. **(b)** Cell viability was assessed by resazurin viability assay before and after cell incubation for 24 h with RA (0.83 mM) or the vehicle control (PBS). Results are expressed as the ratio of the resazurin fluorescence after RA incubation to the fluorescence before incubation. **(c)** To measure invasion, cells pre-treated with RA (0.83 mM) for 2 h were seeded into the upper chamber of Matrigel-coated transwells in serum-free medium. After 24 h, cells that invaded were fixed and stained with DAPI. Scale bar = 100 µm. **(d)** Quantification was performed by counting DAPI-stained cells from 10 randomly-selected fields per insert. Values were normalized to the control, set at 100%. Results are represented as mean ± s.d. of three independent experiments. Statistical significance was determined using one-way ANOVA with Dunnett’s multiple comparison test (a) or unpaired t-test (c) (*P<0.05).

### Rosmarinic acid impairs lysosome degradative capacity

RA has been proposed in the literature to inhibit the P2X7 receptor, leading to reduced intracellular Ca^2+^ levels. To investigate whether this effect could explain why RA impairs lysosome exocytosis, we measured MDA-MB-231 intracellular Ca^2+^ levels, upon incubation with different concentrations of RA, using the Fluo-4 AM probe. Surprisingly, MDA-MB-231 cells showed an increasing trend in intracellular Ca^2+^ levels, although not statistically significant, with increasing concentrations of RA, when compared with the vehicle control (Fig. 4a). This contradicts our initial hypothesis that RA decreases lysosome exocytosis levels by modulating intracellular Ca^2+^ levels through P2X7 inhibition. Therefore, we investigated if lysosome morphology, number and intracellular distribution were affected in MDA-MB-231 cells, upon RA incubation for 24h, by labelling cells with an anti-LAMP1 antibody. Confocal immunofluorescence microscopy showed that there were no detectable differences in LAMP1-positive vesicle morphology, number or distribution in MDA-MB-231 cells incubated with RA when compared with the vehicle control (Fig. 4b). Consistently, western blot analysis showed similar LAMP1 protein levels in RA-incubated and control cells, further suggesting that lysosome number is not affected (Fig. 4c, d). Furthermore, we investigated lysosome acidity using Lysotracker Red DND-99 and found that MDA-MB-231 cells incubated with RA display similar Lysotracker Red DND-99 intensity signal compared with control cells (Fig. 4e), suggesting that lysosome acidity is unaffected by RA. Next, we investigated MDA-MB-231 degradative capacity after incubation with RA. We found the mean fluorescence intensity (MFI) of Magic Red cathepsin L, a cell-permeable substrate that becomes fluorescent when cleaved by active cathepsin L, significantly reduced in MDA-MB-231 cells incubated with RA when compared with control cells (Fig. 4f), suggesting reduced cathepsin L activity. Consistent with these findings, we also observed a significant decrease in the MFI of DQ-Red BSA (Fig. 4g), upon RA treatment, suggesting that RA decreases lysosome degradative capacity despite no detectable differences in lysosome acidity. These observations were further corroborated by western blot analysis of cathepsin D (CTSD), which showed a reduction in the mature and active form of CTSD upon RA incubation (Fig. 4h, i, j). Altogether, these results indicate that RA impairs lysosomal degradative capacity, which may contribute to the decrease in lysosome exocytosis levels observed.

**Figure 4.**
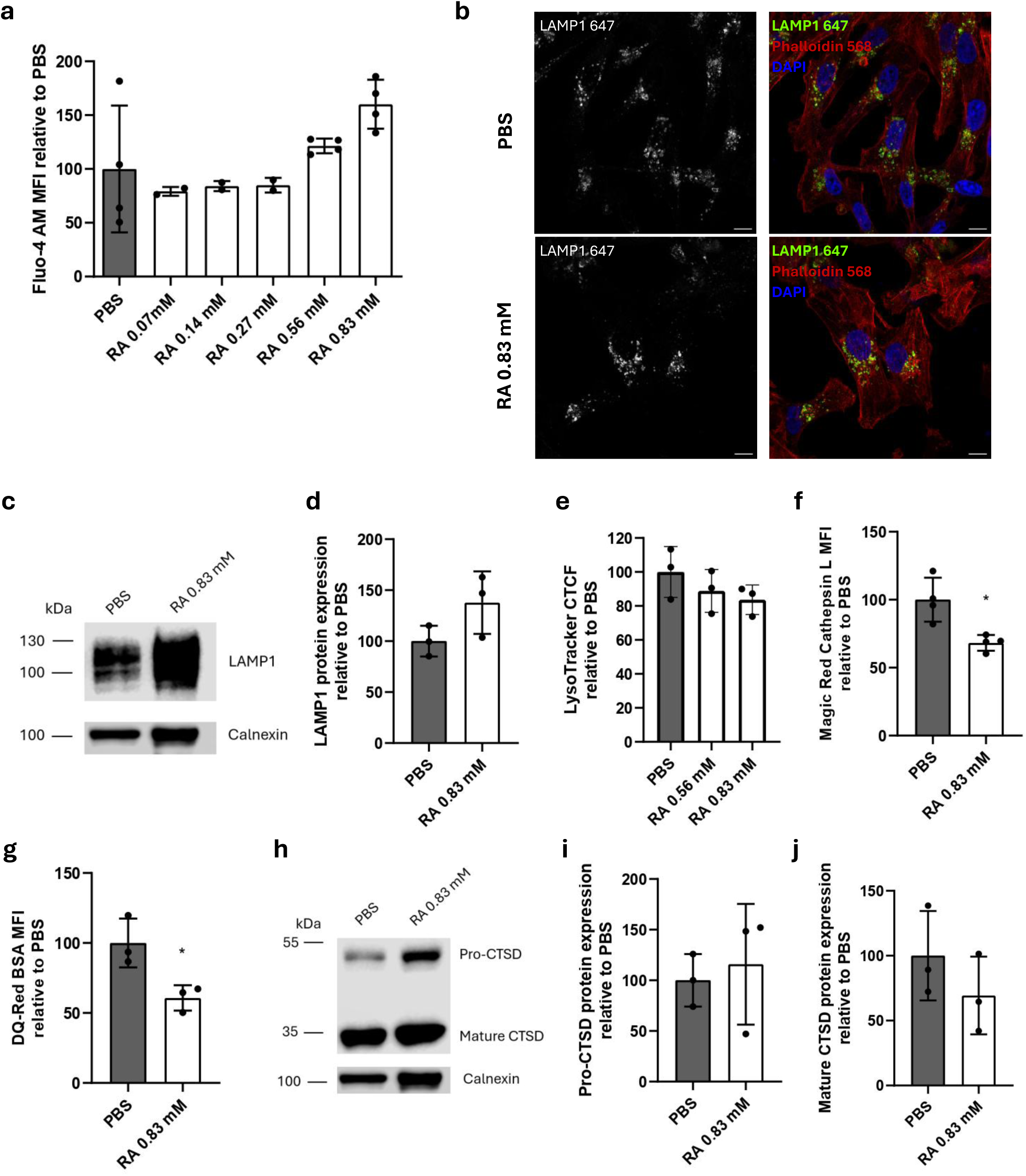
Rosmarinic acid impairs lysosome degradative capacity of triple-negative breast cancer cells. MDA-MB-231 cells were incubated with RA (0.83 mM) or the vehicle control (PBS) for 24 h **(a)** Intracellular Ca^2+^ levels were assessed by measuring the mean fluorescence intensity (MFI) of Fluo-4 AM by flow cytometry. **(b)** Representative confocal images of cells stained with mouse anti-LAMP1 647 (green), phalloidin 568 (red) and DAPI to mark nuclei (blue). Scale bar = 20 μm. **(c)** Western blot with mouse anti-LAMP1 antibody and goat anti-calnexin antibody, used as an housekeeping protein. Original blots are shown in Fig. S4. **(d)** Quantification of total LAMP1 protein levels normalized to calnexin. **(e)** Live cell imaging of MDA-MB-231 cells incubated for 1 h with LysoTracker Red DND-99 (red). Corrected total cell fluorescence (CTCF) was quantified in 50 cells for each condition. **(f)** Cathepsin L activity was assessed by quantifying Magic Red Cathepsin L MFI by flow cytometry. **(g)** Lysosome proteolytic activity was assessed by measuring DQ-Red BSA MFI by flow cytometry. **(h)** Western blot with goat anti-cathepsin D (CTSD) antibody, which detects pro-CTSD and mature CTSD and goat anti-calnexin. Original blots are shown in Fig. S4. **(i)** Quantification of total pro-CTSD protein levels normalized to calnexin. **(j)** Quantification of mature CTSD protein levels normalized to calnexin. Results were normalized to the vehicle control (PBS), in which levels were considered 100%, and are represented as mean ± s.d. of three independent experiments. Statistical significance was determined using one-way ANOVA with Dunnett’s multiple comparison test (a) or unpaired t-test (d, e, f, g, i, j) (*P<0.05).

## Discussion

Lysosome exocytosis is a highly regulated and conserved cellular process that occurs in a wide variety of cell types. In cancer, this mechanism is recognized as a contributor to tumor progression across multiple malignancies ^6,7^. Consistent with this, our findings demonstrate that lysosome exocytosis is significantly enhanced in highly-invasive TNBC cell lines when compared to low-invasive and non-tumorigenic, non-invasive cell lines. This association between invasive behavior and lysosome exocytosis supports a functional role for this pathway in BC progression. Mechanistically, it is described that lysosome exocytosis results in the extracellular release of hydrolytic enzymes, which consequently promotes ECM remodeling and degradation. These changes enable cancer cell invasion and metastatic dissemination ^6–9^. In BC, it was demonstrated that in radiation-surviving BC cells, the peripheral distribution of lysosomes, mediated by ARL8B and the Biogenesis of Lysosome-related Organelles Complex-1-Related Complex (BORC), was associated with increased lysosome exocytosis, increased protease secretion and increased cancer cell motility and invasion. However, this was not shown in non-radiation-treated BC cells ^15^. Furthermore, TRPML1, a Ca^2+^-permeable ion channel located mainly in the membrane of lysosomes and with a relevant role in lysosome exocytosis, when inhibited reduces TNBC cell proliferation, invasion and tumor growth by downregulating the release of lysosomal ATP to the extracellular environment ^28^. However, a clear link between lysosome exocytosis levels and cell invasive phenotype is still lacking. Therefore, we hypothesized that lysosome exocytosis inhibition is sufficient to impair the invasive behavior of TNBC MDA-MB-231 cells. Indeed, we showed that the silencing of RAB10 led to a significant reduction in β-hexosaminidase secretion and impaired the invasive capacity of these TNBC cells. These findings are consistent with previous studies conducted in THP-1 and HeLa cells, reporting that Rab10 silencing impairs lysosome exocytosis ^19,22^, which suggests a conserved role for RAB10 in lysosome exocytosis regulation across different cancer cell lines, including TNBC. Moreover, RAB10 expression has also been associated with tumor growth and poor prognosis in several types of cancer ^29^. In BC cell lines was shown to promote proliferation, migration, and invasion^30^, potentially through the modulation of several signalling pathways, including the Fanconi anaemia pathway, Hippo signalling pathway, and immune-related pathways ^30^. In addition, our findings suggest that RAB10 contributes to TNBC cell invasion, at least in part, through the regulation of lysosome exocytosis. Indeed, we were able to show that RAB10 silencing reduces lysosome exocytosis and gelatin degradation, which mimics the ECM, without affecting invadopodia number. These results suggest that RAB10-dependent lysosome exocytosis contributes to the degradative function of invadopodia rather than to their formation. These findings also support a role for lysosome exocytosis in the release of degradative enzymes, namely cathepsins, to the extracellular space, contributing to ECM degradation and facilitating cancer cell migration and invasion. Although the precise molecular mechanism by which RAB10 regulates lysosome exocytosis remains to be elucidated, our results clearly establish RAB10 as a positive regulator of this pathway and a relevant molecular target to impair TNBC cell invasion.

In addition to genetic modulation, we explored pharmacological strategies to impair lysosome exocytosis as these have greater potential for clinical translation. Surprisingly, neither vacuolin-1 nor verapamil, two known compounds described to modulate lysosome exocytosis in other cells ^6,23,24^, significantly affected lysosome exocytosis in MDA-MB-231 TNBC cells. The lack of effect of these small molecules in TNBC cells suggests a cell type-specific regulation of lysosome exocytosis. Therefore, we searched for other compounds that could potentially impair lysosome exocytosis, namely low molecular weight phenolic compounds (LMWPCs) that target purinergic receptors, known to control intracellular Ca^2+^ levels and exocytosis ^26^. LMWPCs such as RA and resveratrol are hypothesized to modulate the purinergic system, interfering with lysosomal-related pathways ^31,32^. Notably, several of these molecules have been described as anti-metastatic agents in different cancer cell models ^25,26^, further supporting their potential as inhibitors of lysosome exocytosis-driven cell invasion. Therefore, we tested RA in MDA-MB-231 cells and observed a significant decrease in β-hexosaminidase release, in a concentration-dependent manner. Furthermore, we observed that RA impairs MDA-MB-231 cell invasive capacity. Together, these results not only validate our hypothesis but demonstrate that pharmacological inhibition of lysosome exocytosis is sufficient to impair TNBC cell invasive behavior. RA has been proposed to inhibit P2X7 receptor signaling, thereby reducing Ca^2+^ influx, but, surprisingly, we did not observe a reduction in intracellular Ca^2+^ levels following RA incubation. Moreover, we observed that RA does not affect lysosome number, but reduces lysosome degradative capacity. Indeed, altered lysosomal function, including impaired degradative capacity, has been reported along with observation of increased cytosolic Ca^2+^ ^33^. In addition, impaired lysosome degradative function may contribute to the reduction in lysosome exocytosis through defective lysosomal trafficking. For instance, reduced levels of cathepsin B and D or inhibition of lysosomal proteolytic activity were associated with persistent perinuclear lysosome distribution in fibroblasts and slower axonal transport of lysosomes ^34,35^, demonstrating that impaired lysosome degradative activity can lead to defective lysosomal trafficking ^34,35^. Therefore, by interfering with lysosome degradative capacity, RA possibly affects lysosome trafficking to the PM, contributing to the reduced lysosome exocytosis observed.

Collectively, our results reveal not only that more invasive BC cell lines show increased lysosome exocytosis, but also that reducing lysosome exocytosis decreases cell invasive capacity. These findings support lysosome exocytosis as a key driver of cancer cell invasiveness. Through the release of lysosomal degradative enzymes to the extracellular space, it can promote ECM degradation and remodeling, thereby facilitating cancer cell invasion (Fig. 5). By demonstrating that RAB10 silencing or RA-mediated inhibition of lysosome exocytosis significantly impair TNBC cell invasion, we provide experimental evidence that lysosome exocytosis represents a mechanism contributing to the aggressive phenotype observed in these cells and that its targeting is a viable and promising approach to reduce cancer cell aggressiveness. Therefore, our findings open a new avenue for future studies aimed to develop lysosome exocytosis-targeted therapies to attenuate metastatic progression in BC.

**Figure 5.**
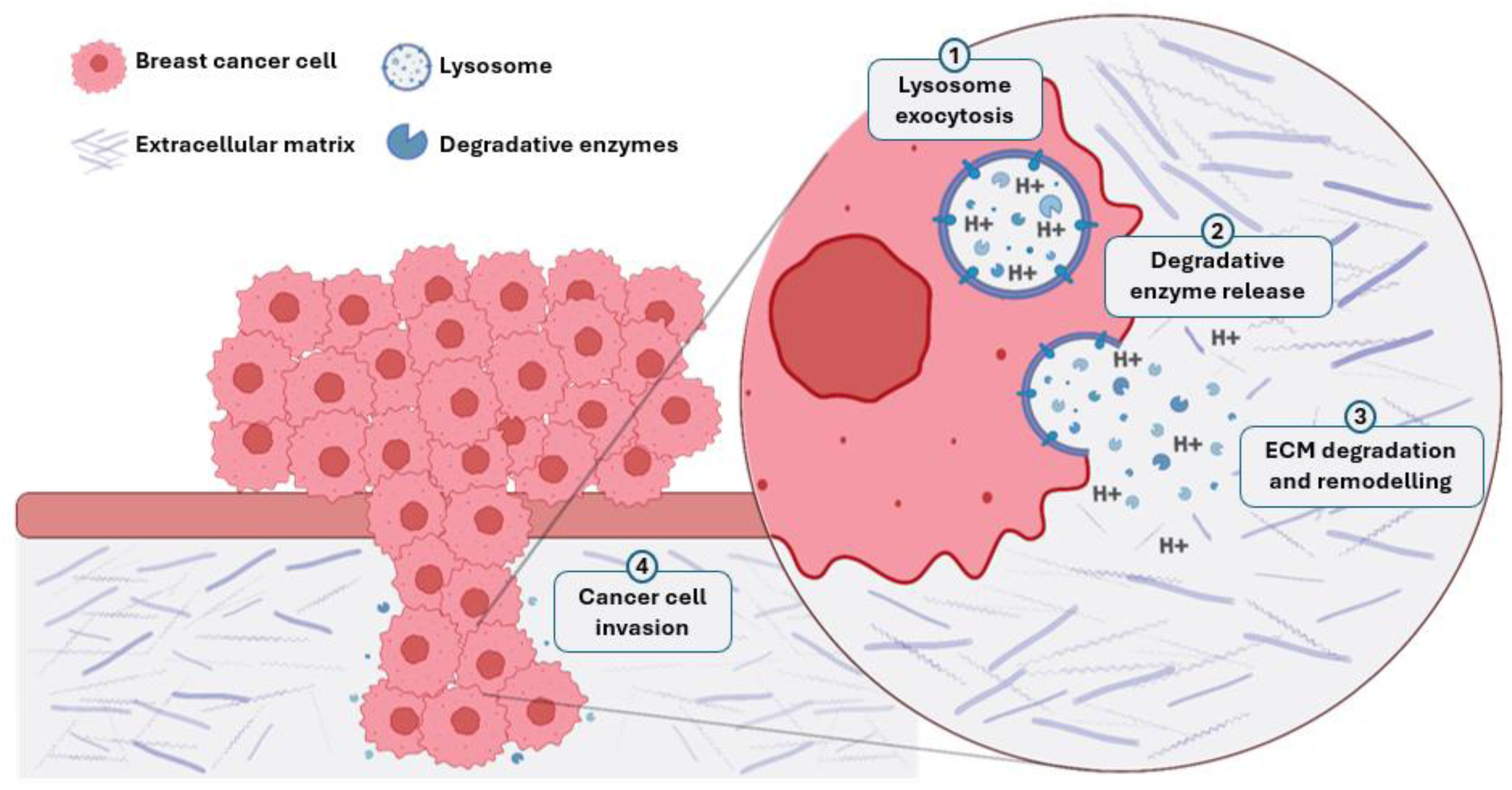
Schematic model of the proposed role of lysosome exocytosis on TNBC cell invasion. BC cells undergo lysosome exocytosis (1), releasing degradative enzymes such as cathepsins to the ECM (2). These enzymes contribute to ECM degradation and remodeling (3), facilitating cancer cell invasion (4). Image created with Inkscape.

## Materials and Methods

### Cell culture

MCF10A cells were a kind gift from Dr. Susana Nunes-Santos (NOVA Medical School, Portugal) and were maintained in high Ca^2+^ medium – Dulbecco’s Modified Eagle Medium/ Nutrient Mixture F-12 (DMEM/F12, Gibco) supplemented with GlutaMAX (Gibco), 5% (v/v) horse serum (Sigma-Aldrich), 10 mM HEPES (Gibco), 1.05 mM CaCl_2_ (Sigma), 100 U/ml penicillin and 100 μg/ml streptomycin (Pen/Strep, GIBCO), 20 ng/mL EGF (Sigma), 10 μg/ml insulin (Gibco), 0.5 μg/ml hydrocortisone (Sigma-Aldrich), and 100 ng/ml cholera toxin (Sigma-Aldrich). MCF-7 and MDA-MB-231, kindly provided by Dr. Gabriela Silva (iBET, Portugal). HCC1806 cells were provided by Dr. Catarina Brito (ITQB, Portugal) were cultured in Roswell Park Memorial Institute (RPMI) complete medium – RPMI (Gibco) supplemented with 10% (v/v) FBS, 0,6% (v/v) HEPES and Pen/Strep. All cell lines were cultured at 37ᵒC and 5% CO_2_, in a humidified incubator.

### Incubation with compounds

MDA-MB-231 at a confluency of ± 80% were incubated with various concentrations of RA (Merck, R4033) (0.07, 0.14, 0.27, 0.56, 0.83 and 1.11 mM) for 24 h, in DMEM without phenol red complete medium – DMEM supplemented with 10 % (v/v) FBS (Gibco), 5 mM GlutaMAX (Gibco), 15 mM HEPES and Pen/Strep. As a control, cells were incubated with the vehicle (PBS).

### Ca^2+^-induced lysosome exocytosis assay

Lysosome exocytosis was induced by incubating cells with 1 µM ionomycin in the presence of 4 mM CaCl_2_ diluted in Ca^2+^- and Mg^2+^-free ice-cold Hanks Balanced Salt Solution (HBSS, Gibco), for 5 min. at 37°C.

### β-hexosaminidase secretion assay

MDA-MB-231 cell supernatants were collected and cells were lysed with 1% IGEPAL. β-hexosaminidase secretion assay was performed as previously described ^18,36^. Briefly, β-hexosaminidase activity was determined by incubating diluted cell supernatants (1:5 in dH_2_O) or lysates in a 96-well plate with 6 mM of the substrate 4-methylumbelliferyl-N-acetyl-β-d-glucosaminide (4-MU-β-D-GlcNAc; Glycosynth) resuspended in substrate buffer (40 mM sodium citrate and 88 mM Na_2_PO_4_, pH 4.5) for 15 min. at 37°C. Fluorescence was measured in a SpectraMax i3X plate reader at 365 nm excitation and 450 nm emission. The protein content from cell supernatants and lysates was determined simultaneously by using the BCA protein assay kit (Pierce Laboratories) as described by the manufacturer. Absorbance was measured at 560 nm in the same plate reader. DMEM complete medium diluted 1:5 and 1% IGEPAL was used as a control. β-hexosaminidase (β-hex) activity was calculated for each sample normalizing to total protein amount as following: β-hex activity in supernatant=(fluorescence (365/450)−HBSS alone)/protein µg. β-hex activity in cell lysate=(fluorescence (365/450)−IGEPAL alone)/protein µg. Total β-hex activity=β-hex activity in supernatant+5× β-hex activity in cell lysate. Finally, the percentage of β-hex release was calculated as following: β-hex release (% of total)=100×(β-hex activity in the supernatant/total β-hex activity).

### Transwell invasion assay

MDA-MB-231 cells incubated overnight (ON) in serum-free DMEM supplemented with 0.5% bovine serum albumin (BSA) were placed on the top of 8 μm pore transwells coated with Matrigel (Corning). Cells treated with RA, were first incubated with the compounds for 2 h before being seeded in the transwells. Complete medium was added to the lower chamber to induce cell migration and invasion, and cells were allowed to invade for 24 h at 37°C, 5% CO_2_. Medium was removed from the upper chamber and cells that invaded into the lower chamber were fixed with 4% (v/v) paraformaldehyde (PFA; Alfa Aesar) and nuclei stained with 1 μg/ml of DAPI. Ten randomly-selected fields of each transwell were acquired on an Axiovert 40 (Zeiss) microscope using a digital camera (AxioCam MRc) and ZEN 2 software (blue edition) and analyzed with ImageJ software 1.8.0.

### Gelatin degradation assay

Coverslips were coated with 50 μg/ml of poly-D-Lysine for 1 h and incubated with 0.5% glutaraldehyde for 15 min. After incubation with 0.2 mg/ml of Oregon Green ®-conjugated pig skin gelatin (Invitrogen), diluted in PBS with 0.2 % sucrose for 10 min., cells were seeded and allowed to adhere for 6 h at 37°C, 5% CO_2_. Cells were fixed for 15 min. in 4% paraformaldehyde (PFA) (Alfa Aesar) and blocked/permeabilized for 30 min. in PBS containing 0.5% BSA and 0.1% saponin. Cells were then incubated with primary antibody mouse anti-cortactin antibody – clone 4F11 (Millipore) (1:300) – followed by incubation with Alexa Fluor 568-conjugated goat anti-mouse secondary antibody (Invitrogen) (1:1000). Next, coverslips were mounted with Fluoromount-G with DAPI (Invitrogen). Images were acquired using a Zeiss LSM 710 confocal microscope equipped with a Plan Apochromat 63/1.40 Oil Ph3 lens and Zen Blue 2010b SP1 software and image analysis was performed on ImageJ. For each independent experiment, 20 random fields were imaged. Percentage of gelatin degradation was quantified by calculating the area of the dark spots in the Oregon Green-labeled gelatin that overlaps with cells, over the total area of the image, multiplied by 100. Active invadopodia were assessed by colocalization between cortactin-positive structures and areas of gelatin degradation. The percentage of cells with active invadopodia was calculated as the number of active invadopodia-positive cells divided by the total number of cells in the field, multiplied by 100.

### Resazurin viability assay

MDA-MB-231 cells (2.5 x 10^5^) were seeded in 96-well plates. On the following day, resazurin 1x (Invitrogen) diluted in DMEM complete medium without phenol red was added to each well and incubated for 1 hour 20 min. at 37°C. Fluorescence was measured in a SpectraMax i3X plate reader at 531 nm excitation and 595 nm emission. RA was then added to the cells at different concentrations (0.07, 0.14, 0.27, 0.56, 0.83 mM) in DMEM complete medium without phenol red. Twenty-four h after incubation,resazurin 1x was added again to the cells and measured as described above. Resazurin 1x fluorescence in the absence of cells was used as the background. Relative cell viability was calculated by dividing the background corrected fluorescence values measured after RA incubation by the corresponding fluorescence values measured before incubation.

### Flow cytometry

MDA-MB-231 cells incubated for 24 h with RA (0.83 mM) or the vehicle (PBS) were stained with an Alexa Fluor 488 anti-LAMP1 antibody, clone H4A3 (BioLegend) (1:1000), for 30 min. at 4°C, washed and resupended in PBS, 1% FBS and 2 mM EDTA. Immediately before acquisition, propidium iodide (PI) was added at a final concentration of 0.5 μg/ml to exclude non-viable cells.

To assess intracellular Ca^2+^ levels, cells were incubated with 5 μM of Fluo-4AM (Molecular Probes), for 30 min. at 37°C. After washing with PBS, cells were resuspended in PBS, 1% FBS and 2 mM EDTA and incubated for 30 min. to allow de-esterificaton of the intracellular probe. To analyze degradative capacity, cells were incubated with 10 μg/ml DQ-Red BSA (Invitrogen) ON, or Magic Red Cathepsin L 1x (BioRad), for 1 h. Upon incubation, cells were washed in PBS and resuspended in PBS with 1% FBS and 2 mM EDTA. Acquisition was performed in a FACS CANTO II (BD Biosciences) flow cytometer and at least 30,000 cells were analyzed using FlowJo version 10.1r7 software.

### Immunofluorescence

MDA-MB-231 cells were grown ON in coverslips. On the following day, cells were incubated with RA (0.83 mM) or the vehicle (PBS) in DMEM complete medium without phenol red. After 24 h, cells were fixed for 20 min. at RT with 4% paraformaldehyde (PFA) and blocked and permeabilized in PBS containing 0.5% BSA and 0.1% saponin for 30 min., at RT. Next, cells were incubated with primary antibody, namely anti-human Alexa Fluor 647 LAMP1 (BioLegend) (1:500) and Alexa Fluor 568 Phalloidin (Invitrogen) (1:200), for 1 h at RT. Lastly, cells were mounted in Fluoromout G (Invitrogen) mounting media with DAPI. Image acquisition was performed using a Zeiss LSM710 confocal microscope, equipped with a 63x 1.4 NA oil objective. Image processing and analysis was done using FIJI software version 2.9.0.

### Live cell imaging

MDA-MB-231 cells were seeded in Nunc™ Lab-Tek™ Chambered Coverglasses (Thermo Fisher Scientific) and allowed to adhere ON. On the following day, cells were incubated with RA (0.83 mM) or the vehicle (PBS) for 24 h. After incubation with 10 μM LysoTracker Red DND-99 (Invitrogen) DMEM complete medium without phenol red, cells were further incubated for 15 min with 0.2 μg/ml Hoescht 33342 (BioRad) (1:1000) in DMEM complete medium without phenol red. After extensive washing with PBS, cells were kept in medium. Live-cell imaging was performed using a Zeiss LSM 710 confocal microscope equipped with a Plan Apochromat 63/1.40 Oil Ph3 lens and Zen Blue 2010b SP1 software and image analysis was performed on FIJI software version 2.9.0.

### Immunoblotting

MDA-MB-231 cells previously incubated with RA (0.83 mM) or the vehicle (PBS) were lysed for 30 min. on ice, in lysis buffer (50 mM Tris–HCl pH 7.5, 1 mM EDTA, 1 mM EGTA, 150 mM NaCl, 2 mM MgCl2, 1 mM DTT, 0.1% IGEPAL), supplemented with 25% protease inhibitors cocktail (Roche) and 100 μM orthovanadate (Sigma), followed by centrifugation at 14,000 x *g* for 30 min. at 4°C. The supernatant was collected, and protein quantification was performed using the DC Protein Assay Kit (Bio-Rad Laboratories), according to the manufacturer’s instructions.

Thirty µg of total protein were loaded on a 10% SDS-polyacrylamide gel, transferred to a nitrocellulose membrane (GE Healthcare Life Sciences), for 1 h at 100 V and processed for western blotting.

Membranes were blocked for 1 h, at RT, using a blocking solution (5% BSA in TBS with 0,1% Tween-20). Membranes were then incubated ON, at 4°C with primary antibodies rabbit anti-human LAMP1 (Cell Signalling) (1:1000) or goat anti-human CTSD (Sicgen) (1:1000). On the following day, membranes were incubated, for 1 h, with horseradish peroxidase (HRP)-conjugated secondary antibodies goat anti-rabbit (BioRad) (1:7500) or donkey anti-goat (ThermoFisher Scientific) (1:7500). Membrane stripping was performed by incubating the membrane in stripping buffer (2% SDS, 100 mM β-mercaptoethanol, 62.5 mM Tris-HCl pH 6.7), for 30 min. at 55°C, followed by re-incubation with blocking solution. Stripped membranes were incubated with primary antibody goat anti-human calnexin 1 (Sicgen) (1:1000), as previously described, followed by incubation with HRP-conjugated secondary donkey anti-goat antibody. HRP activity was detected using Amersham ECL Prime (GE Healthcare Life Sciences), according to the manufacturer’s instructions, in a LICOR Odyssey FC (LI-COR Biosciences, Lincoln, NE, USA) imaging system. Bands were quantified using FIJI software version 2.9.0.

### Statistical analysis

Numerical data are presented as mean±standard deviation (±s.d.). One-way ANOVA followed by Dunnett’s multiple comparisons test was used to compare different data sets with a control. Unpaired t-tests were employed to compare the means between two independent data sets. Statistical analysis was performed using GraphPad Prism version 8.0.2.

## Acknowledgments

We would like to thank the NMS microscopy, flow cytometry and cell culture core facilities for the scientific and technical assistance, as well as Dr. Susana Nunes-Santos (NOVA Medical School, Portugal) for the kind gift of MCF10A cells, Dr. Gabriela Silva (iBET, Portugal) for MCF-7 and MDA-MB-231 cells and Dr. Catarina Brito (ITQB, Portugal) for HCC1806 cells.

## Author contributions

A.R.R., C.E., T.B. and D.C.B. conceptualized and designed the study. A.R.R., C.E., I.F., M.F., and M.G.O. acquired the experimental data. J.S.R. provided resources and contributed to the methodology employed. A.R.R., C.E., D.C.B. analyzed and interpreted the data and drafted the initial manuscript. C.E., T.B. and D.C.B. supervised the study and D.C.B provided funding. All authors reviewed the final version of the manuscript and approved its submission.

## Data availability

All relevant data can be found within the article and its supplementary information.

