## SupplementaryInformation for "Lysosome Exocytosis Inhibition Impairs Triple-Negative Breast Cancer Cell Invasion"

[1] iNOVA4Health, NOVA Medical School, Faculdade de Ciências Médicas, Universidade NOVA de Lisboa, Campo dos Mártires da Pátria, 130, Lisboa 1169-056, Portugal.

[#] These authors contributed equally to this work

* Correspondence: Duarte C. Barral, NOVA Medical School, Faculdade de Ciências Médicas, Universidade NOVA de Lisboa, Campo dos Mártires da Pátria, 130, Lisboa 1169-056, Portugal.

**Supplementary Figure S1**

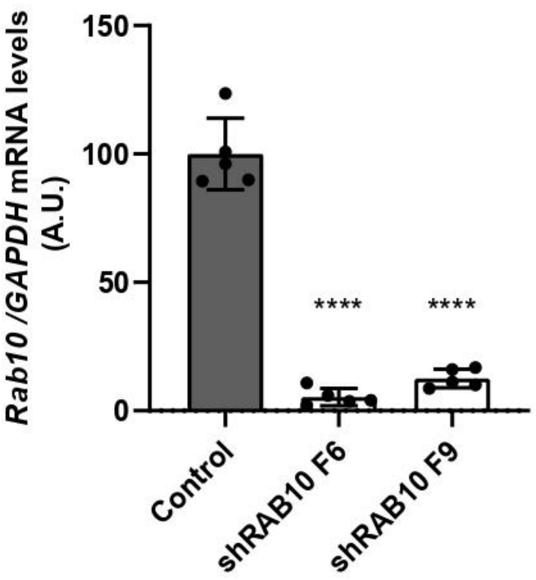

**Supplementary Figure S2**

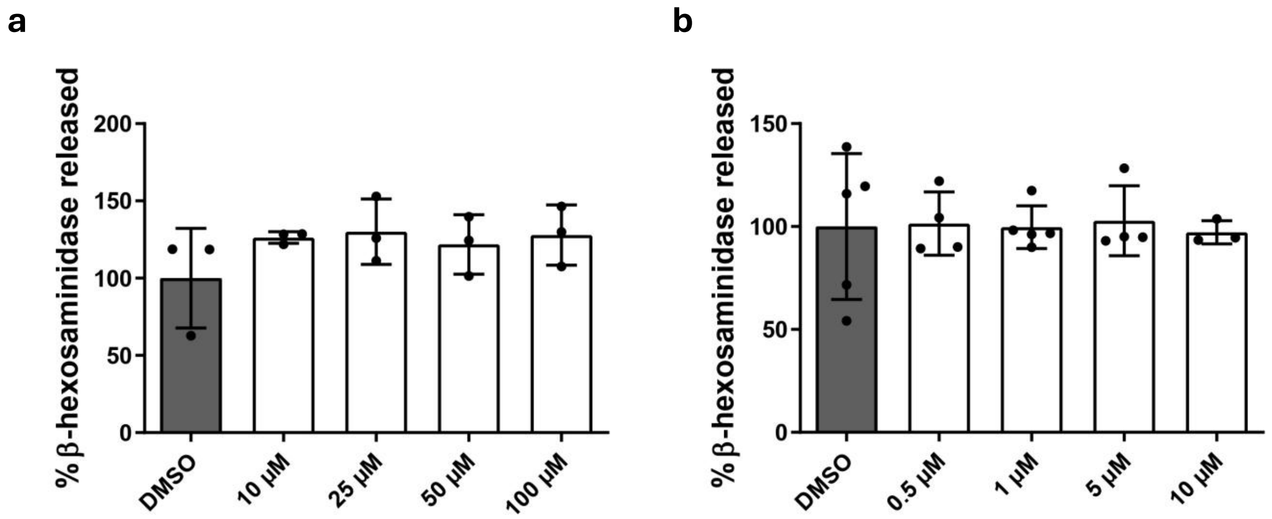

**Supplementary Figure S3**

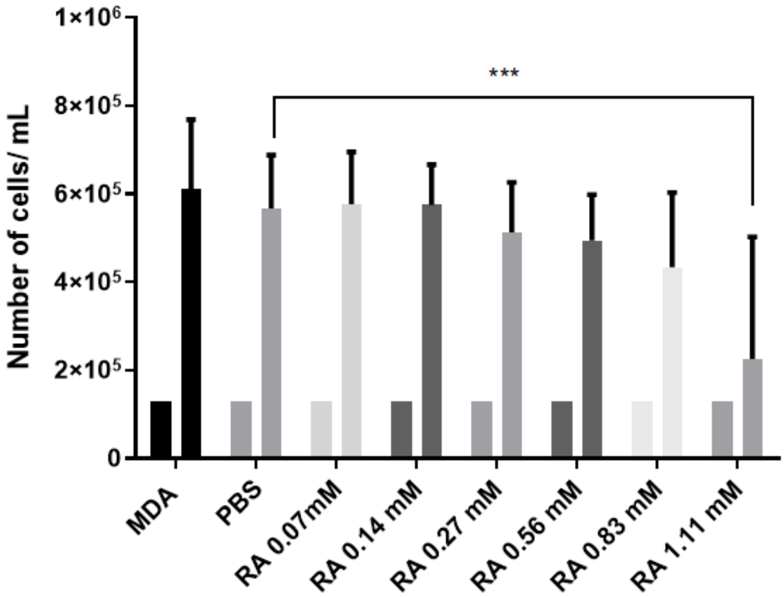

**Supplementary Figure S4**

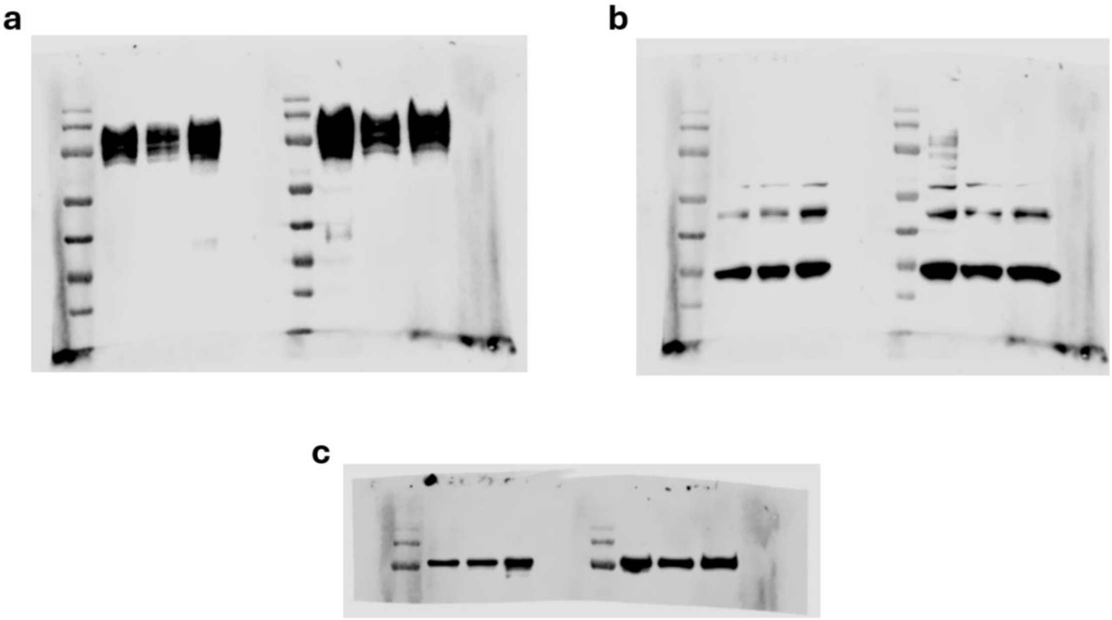

**Supplementary Figure legends**

**Supplementary Figure S1 – RAB10 silencing in triple-negative breast cancer cells.** MDA-MB-231 cells were transduced with lentiviruses encoding shRNA hairpins targeting RAB10 (shRAB10 F6 and F9), and a non-targeting shRNA (Mission) as a control. Silencing efficiency was determined by RT-qPCR, measuring mRNA expression levels of RAB10, relative to the housekeeping gene GAPDH. Statistical significance was determined using one-way ANOVA with Dunnett's multiple comparison test (****$P<0.0001$). A.U., arbitrary units.

**Supplementary Figure S2 – Verapamil and vacuolin-1 do not affect lysosome exocytosis in triple-negative breast cancer cells.** MDA-MB-231 cells were incubated with **(a)** verapamil hydrochloride (Merck, V4629) at different concentrations (10, 25, 50 and 100 μM), or **(b)** vacuolin-1 (Santa Cruz, sc216045) at different concentrations (0.5, 1, 5 and 10 μM), in DMEM complete medium, for 24 h, with DMSO as the vehicle control. β-hexosaminidase release was measured in cell supernatants and extracts and quantified as described in the Materials and Methods. Results are represented as mean ± s.d. of at least three independent experiments. Statistical significance was determined using one-way ANOVA with Dunnett's multiple comparison test.

**Supplementary Figure S3 – Rosmarinic acid does not affect MDA-MB-231 cell proliferation for concentrations up to 0.83 mM.** MDA-MB-231 ($1x10^5$) cells were plated and let to adhere overnight. On the next day, cells were incubated with different concentrations of rosmarinic acid (0.07, 0.14, 0.27, 0.56, 0.83 and 1.11 mM), or the vehicle control (PBS), in DMEM complete medium without phenol red. After 24 h, cells were collected and viable cells counted using trypan blue to exclude dead cells. Results are presented as number of viable cells/ml before (day 1) and after incubation with rosmarinic acid (day 3). Results are represented as mean ± s.d. of three independent experiments. Statistical significance was determined using one-way ANOVA with Dunnett's multiple comparison test (***$P<0.001$).

**Supplementary Figure S4 – Uncropped western blots.** **(a)** Incubated with primary antibody rabbit anti-human LAMP1 and HRP-conjugated goat anti-rabbit secondary antibody. **(b)** Incubated with primary antibody goat anti-human CTSD that labels pro-CTSD and mature CTSD and HRP-conjugated donkey anti-goat secondary antibody. **(c)** Incubated with primary antibody goat anti-human calnexin 1 and HRP-conjugated donkey anti-goat secondary antibody.

**Supplementary Tables**

**Supplementary Table S1** - shRNAs used to transduce MDA-MB-231 cells .

| Gene | Association Number | Hairpin | Source/ Clone ID | Target sequence (5'-3') |
|---|---|---|---|---|
| RAB10 | NM_016131 | F6 | TRCN0000029189 | GCCTTCAATACTACCTTTATT |
| | NM_016131 | F9 | TRCN0000029193 | CAAGTGTGATATGGACGACAA |

**Supplementary Table S2 -** Primers used for qRT-PCR.

| Gene | Primer Forward (5'-3') | Primer Reverse (5'-3') |
|---|---|---|
| *GAPDH* | CATTTCCTGGTATGACAACGA | GTCTACATGGCAACTGTGAG |
| *Rab10* | GTGTATGACATCACCAACGG | GCAGCACTTGCTCTTCCAGC |

**Supplementary Materials and Methods**

**Cell culture**

HEK-Star-Rdpro cells (obtained from Culture Collections, UK Health Security Agency, Porton Down, Salisbury, United Kingdom) were maintained in DMEM complete medium – DMEM supplemented with 10 % (v/v) fetal bovine serum (FBS, Gibco), 5 mM GlutaMAX (Gibco), 15 mM HEPES and Pen/Strep.

**shRNA transduction**

Lentiviruses were produced in HEK-Star-Rdpro cells in accordance with BL-2+ protocols publicly available at the Broad Institute's RNAi Consortium website (https://portals.broadinstitute.org/gpp/public/resources/protocols). Packaging (psPAX/GAG) and envelope plasmids, PMD2G (VSV-G) and Hairpin-pLKO.1 vector were obtained from the RNAi Consortium. The shRNA sequences are listed in Supplementary Table S1. As a control, a non-targeting sequence (Mission; 5'-CAACAAGATGAAGAGCACCAA-3'; Sigma) was used.

For lentiviral transduction, MDA-MB-231 cells plated in 6-well plates were incubated with shRNA-encoding lentivirus supernatants in the presence of 8 μg/ml polybrene (hexadimethrine bromide; Sigma). Twenty-four hours after infection, virus-containing medium was replaced with selection medium containing 1 μg/ml puromycin (Sigma). All assays were performed 7–8 days after selection.

**RNA isolation and qRT-PCR**

Total RNA was isolated with RNeasy Mini Kit (Qiagen), according to the manufacturer's instructions. Synthesis of complementary DNA (cDNA) was performed by incubating 1 μg of total RNA with 1 mM of dNTP mix (Thermo Fisher Scientific) and 0.3 μg/μl of random primers p(DN) (Roche) at 65°C for 5 min., followed by incubation with 5x first strand buffer (Invitrogen), 20 mM DTT (Invitrogen) and 40 U/μl of Recombinant Ribonuclease Inhibitor RNaseOUT (Invitrogen) at 25°C for 2 min. Finally, 50 U/μl of Superscript II Reverse Transcriptase (Invitrogen) was added, samples were incubated at 25°C for 10 min., 42°C for 50 min. and, finally, at 70°C for 15 min. For Real-Time quantitative Polymerase Chain

Reaction (RT-qPCR), FastStart Essential DNA Green Master kit (Roche) was used according to manufacturer's instructions and samples were run and analyzed in a QuantStudio™ 5 Real-Time PCR System from ThermoFisher with the following protocol: pre-incubation at 95°C for 10 min.; 45 amplification cycles at 95°C for 10 sec., 60°C for 10 sec., and 72°C for 20 sec.; melting at 95°C for 10 sec., 65°C for 1 min. and 97°C for 1 sec. Gene expression was calculated relative to the housekeeping gene, GAPDH. The sequences of the primers used are listed in Supplementary Table S2.

**Incubation with compounds**

MDA-MB-231 at a confluency of ± 80% were incubated with various concentrations verapamil hydrochloride (Merck, V4629) at different concentrations (10, 25, 50 and 100 µM), or vacuolin-1 (Santa Cruz, sc216045) at different concentrations (0.5, 1, 5 and 10 µM), in DMEM complete medium, for 24 h, with PBS as the vehicle control, in DMEM complete medium without phenol red – DMEM supplemented with 10 % (v/v) FBS (Gibco), 5 mM GlutaMAX (Gibco), 15 mM HEPES and Pen/Strep. As a control, cells were incubated with the vehicle (PBS).

**Trypan-Blue exclusion**

MDA-MB-231 cells (1.3 x $10^5$) were seeded in 24-well plates and allowed to adhere ON. On the following day, cells were incubated with different concentrations of RA (0.07, 0.14, 0.27, 0.56, 0.83 and 1.11 mM) or the vehicle control (PBS) in DMEM complete medium without phenol red for 24 h. The next day, cells from each condition were collected and diluted in trypan blue. The number of viable cells in each condition was assessed by counting trypan blue negative cells.
